# Timing is everything: event-related transcranial direct current stimulation improves motor adaptation

**DOI:** 10.1101/2021.11.26.470091

**Authors:** Matthew Weightman, John-Stuart Brittain, Alison Hall, R. Chris Miall, Ned Jenkinson

**Affiliations:** School of Sport, Exercise and Rehabilitation Sciences, University of Birmingham, Edgbaston, Birmingham, B15 2TT, UK; School of Psychology, University of Birmingham, Edgbaston, Birmingham, B15 2TT, UK; MRC-Versus Arthritis Centre for Musculoskeletal Ageing Research, University of Birmingham, Edgbaston, Birmingham, B15 2TT, UK; Centre for Human Brain Health, University of Birmingham, Edgbaston, Birmingham, B15 2TT, UK

**Keywords:** TDCS, Motor Adaptation, Plasticity

## Abstract

**Background:** There is a current discord between the foundational theories underpinning motor learning and how we currently apply transcranial direct current stimulation (TDCS): the former is dependent on tight coupling of events while the latter is conducted with very low temporal resolution.

**Objective:** Here we aimed to investigate the temporal specificity of stimulation by applying TDCS in short epochs, and coincidentally with movement, during a motor adaptation task.

**Methods:** Participants simultaneously adapted a reaching movement to two opposing velocity-dependent force-fields (clockwise and counter-clockwise), distinguished by a contextual leftward or rightward shift in the task display and cursor location respectively. Brief bouts (< 3 second) of event-related TDCS (er-TDCS) were applied over M1 or the cerebellum during movements for only one of these learning contexts.

**Results:** We show that when short duration stimulation is applied to the cerebellum and yoked to movement, only those reaching movements performed simultaneously with stimulation are selectively enhanced, whilst similar and interleaved movements are left unaffected. We found no evidence of improved adaptation following M1 er-TDCS, as participants displayed equivalent levels of error during both stimulated and unstimulated movements. Similarly, participants in the sham stimulation group adapted comparably during left and right-shift trials.

**Conclusions:** It is proposed that the coupling of cerebellar stimulation and movement influences timing-dependent (i.e., Hebbian-like) mechanisms of plasticity to facilitate enhanced learning in the stimulated context.

## Introduction

Coincident, time-dependent mechanisms of synaptic plasticity are the canonical basis of theories of motor learning [1, 2]. These ‘Hebbian’ mechanisms are ubiquitous throughout the mammalian brain, having been described in the hippocampus, cerebellum, and sensory-motor cortices [3–5], and are believed to underpin all forms of learning and memory. Yet, protocols for non-invasive brain stimulation intended to promote motor learning and rehabilitation - particularly transcranial direct current stimulation (TDCS) - largely ignore timing-dependent mechanisms.

TDCS is a non-invasive form of brain stimulation often used to induce plasticity in the motor system by modulating neural excitability. Studies measuring the physiological effects of TDCS using transcranial magnetic stimulation (TMS), electroencephalogram (EEG), functional MRI (fMRI), magnetic resonance spectroscopy (MRS) etc. have reported alterations in neural excitability that are dependent on the polarity of stimulation [6–11]. Changes in neuronal excitation have been shown to be almost instantaneous in terms of increases in firing rates [12–14] and motor evoked potentials [6, 15] following a brief application of TDCS and other forms of polarising currents. Despite this, most conventional studies apply TDCS for 15-20 minutes in a continuous stimulation period, prior to and/or during a motor task and these changes in neural excitability do not always reliably translate into modulation behaviour, including motor learning. One potential reason for these apparently inconsistent results may be the low temporal resolution of stimulation. If TDCS can instantaneously modulate neural activity, applying short duration epochs of TDCS temporally aligned with movement has the potential to specifically and selectively enhance learning, by driving coincident mechanisms of plasticity in the circuits of the brain that are active during the movement.

Here we demonstrate (and replicate) that brief epochs of stimulation, applied in synchrony with movement, selectively enhanced motor adaptation whilst, importantly leaving adaptation of the non-stimulated (yet interleaved) movements unaffected.We believe this first proof-of-principle study using a novel stimulation protocol harnesses mechanisms of Hebbian plasticity, resulting in the selective, transient, potentiation of those neurobehavioural circuits that are active concomitant with stimulation [16–18].

## Materials & Methods

### Participant Details

A total of seventy-eight participants (aged 18-32 years, mean = 20.6 ± 3.0 years; 39 male) gave written informed consent to take part in the study (approved by the Science, Technology, Engineering and Mathematics Ethical Review Committee at the University of Birmingham). All participants were right-handed, had normal or corrected to normal vision and completed a safety screening questionnaire for TMS and TDCS prior to beginning the session. For the main experiment sixty participants were pseudo-randomised into one of three experimental groups: M1 er-TDCS group (n = 20; mean age = 19.7 ± 0.8 years, 10 males), Cerebellar er-TDCS group (n = 20; mean age = 19.8 ± 1.9 years, 11 males) or a Sham TDCS group (n = 20; mean age = 19.5 ± 0.8 years, 9 males). A further eighteen naive participants (n = 18, mean age = 23.9 ± 4.7 years, 9 males) were recruited for a secondary experimental group. Participants in this secondary group also received cerebellar er-TDCS. Importantly, data from the secondary group were collected after the conclusion of the main experiment in order to replicate the main experimental finding. No a priori statistical methods were used to determine sample size. Instead, our sample size was chosen to be consistent with, or greater than similar existing literature [19-21].

### Experimental Design

Participants were seated in an armless chair so they could comfortably reach and manipulate the handle of a custom-built robotic manipulandum (vBOT; [22]) with their right arm. The vBOT measured and stored the position and velocity of the handle at 1000Hz, only allowing movements in the horizontal plane. The visual display screen (Mac Cinema HD Display) was reflected in a horizontal mirror (60 × 76 cm) to appear as a virtual image co-planar with the manipulandum. The screen displayed two grey circular markers (2 cm diameter) which represented the home position and the target, located 20 cm and 10 cm away from the edge of the screen respectively. The screen also displayed a white cursor (1 cm diameter) which showed the position of the vBOT handle. The home and target markers were displaced 10cm left or right of the screen midline, depending on the context of the trial, and the cursor was displaced 10cm left or right of the vBOT handle (Figure 1a).

**Figure 1:**
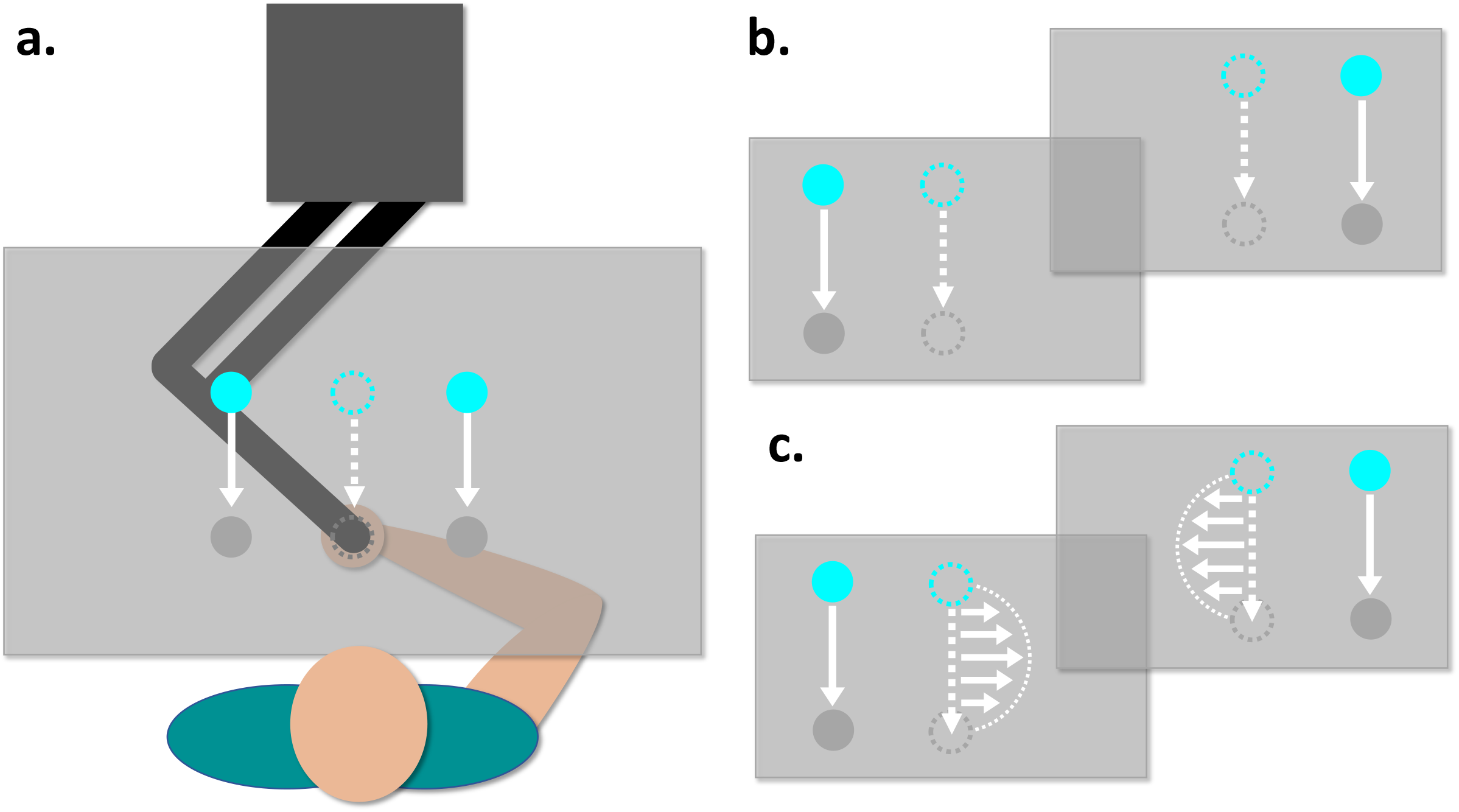
Experimental design and set-up. **a**. A schematic of the task set-up, with an example screen display. Movements were always made in the midline position, but the cursor, home and target markers would be shifted either 10cm to the right or left. The order of the contextual shift in the task display was pseudorandomised, with an equal number in each task phase and runs of no more than three trials of the same contextual shift. Examples of the task display during null field **(b)** and context-dependent force-field **(c)** trials for both trial types.

Participants were told that the aim of the task was to make fast movements from the home marker to the target marker, so that the cursor moved in a straight line between the two (10 cm movement). At the beginning of each trial the participant entered the home position and were held there for 3 seconds by stiff spring forces on the vBOT handle, before being allowed to move. The home marker then changed from grey to blue indicating that participants were allowed to make their movement and the holding forces were released. During the hold period participants were asked to keep relaxed and not to ‘pull’ or ‘lean’ on the handle. To encourage fully completed movements, participants were told that it was acceptable to overshoot the target slightly, but were discouraged from making excessively large movements. If the movement was accurate and hit the target marker it would flash yellow for 1 second indicating a successful trial. If the movement failed to hit the target or deviated ± 2 cm from the midline at any point during the movement path the target would flash red for 1 second, indicating an unsuccessful trial. Once the movement was completed, the vBOT would actively guide the handle back to the home position with a spring force, ready for the next trial to start. Each trial took roughly 5 seconds from start to finish (3 second hold, 1 second movement, 1 second return). Vision of the upper arm was blocked during the task using a curtain and all lights extinguished prior to starting the task.

### Behavioural Protocol

The behavioural task consisted of 600 trials and was split into three phases: Baseline (100 trials), Adaptation (400 trials) and Washout (100 trials). On each trial, in all three phases, the task display would either be presented with a 10 cm leftward or rightward shift from the midline position. The order of this shift was pseudo-randomised, so that there was an equal number of leftward and rightward shift trials in each phase and no more than 3 consecutive trials with the same contextual shift. The trial/shift order was the same for all participants. During baseline trials no forces were imposed on the handle so participants could move between the home marker and the target unperturbed (Figure 1b). For adaptation trials the leftward or rightward contextual shift in the task display was consistently associated with either a clockwise (CW, for left-shift) or counter-clockwise (CCW, for right-shift) velocity sensitive curl force-field (Figure 1c) -thus creating two distinct trial contexts [23]. The strength of both imposed force-fields was 12 N/m/s (see equation 1).

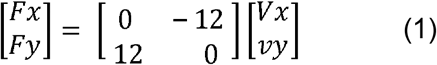

Washout trials immediately followed the adaptation phase, which were once again performed without any forces i.e. identical to baseline. Participants were not given any explicit information regarding the link between the visual shift and associated direction of force-field.

In order to measure compensatory forces applied against the curl field, 60 error-clamp trials were pseudorandomly interleaved throughout the task (10% of all trials). These trials were distributed proportionally throughout the three task phases. During error-clamp trials, movements were constrained to a ‘virtual channel’ between the home position and target marker, so that forces produced against the channel walls could be measured (see Supplementary Figures 2 & 3 for data and analysis). Error-clamp trials took place twice every 20 trials and the contextual shift was pseudorandomly ordered so that no more than two successive error-clamp trials were of the same shift. Additionally, error-clamp trials only occurred on trials where there was a switch in context and not after one or two trials of the same shift. To avoid directional feedback, the vBOT position was indicated via an expanding semi-circle representing only the distance from start location [24]. At the end of the task each participant was asked if they had noticed the relationship between the visuospatial context and direction of the force in order to understand their explicit knowledge during the task.

### Transcranial Direct Current Stimulation

Anodal TDCS was delivered via two sponge electrodes (5 × 7 cm) soaked in saline solution, using a nurostym tES device (Neuro Device Group S.A., Poland). For cerebellar stimulation the anodal electrode was placed over the right cerebellar cortex (3 cm lateral to the inion; [25]) and the cathode was positioned on the superior aspect of the right trapezius muscle [19, 21, 26]. For M1 stimulation the anodal electrode was positioned over the hand area of the left motor cortex, identified by single pulse TMS (Magstim Rapid2 stimulator; Magstim Ltd, UK) delivered at suprathreshold stimulus intensity so as to elicit a visible twitch of the first dorsal interosseous muscle. The cathode electrode was placed on the skin over the contralateral supraorbital ridge [6]. TDCS was only delivered during the adaptation phase, the parameters of which depended on the stimulation group. Once the task had ended, participants rated their perceived comfort and confidence in their belief that they received real stimulation on a 10-point visual analogue scale (VAS). They were also asked if they noticed anything specific regarding the timing of the stimulation, with respect to the task.

In the main experiment, participants in the M1 and cerebellar stimulation groups received brief epochs of TDCS during the adaptation phase, with each epoch, temporally overlapping with movements made through the CCW force-field, when the task display was shifted to the right. Stimulation was ramped up over 1 second during the hold period (1 second prior to the movement cue). It was then held at 2 mA for 1 second during the movement and then ramped down over 1 second as the vBOT returned the participant’s hand to the home position. TDCS was applied on both force-field and error-clamp trials. In the secondary experimental group, the er-TDCS protocol was conducted as per the main experiment, however, stimulation was only applied over the cerebellum and concurrently with movements made during the CW force-field and associated left-shift in visual display. Data from this group served as an attempt to replicate the effect found in the main experiment, with stimulation applied during movements through the opposing force-field direction. For the sham group, stimulation was ramped up over 10 seconds at the start of the adaptation phase, held at 2 mA for 10 seconds and then ramped down over a further 10 seconds. The electrode montage was randomly assigned to either M1 or cerebellar prior to starting the session.

### Data & Statistical Analysis

Data collected from the vBOT were analysed offline in MATLAB (The Mathworks, version R2018b). The lateral deviation (LD) of movements at peak velocity was calculated for null and force-field trials and subsequently averaged into bins. For error-clamp trials, a force compensation ratio (FC) was calculated, see equation 2.

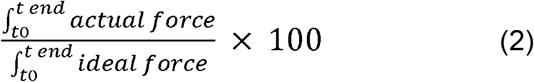

Actual force was the forces generated against the channel walls, integrated across the movement; ideal force was the force required to fully compensate for the perturbation in force-field trials (ideal force: velocity x field constant). Any LD or FC values that fell outside ± 2 SD of the mean across the group were excluded prior to averaging and thus removed from further analysis. LD and FC values were analysed separately for the two contexts. Area under the learning curve for LD and FC during each phase of null, force-field and error-clamp trials was calculated to be used for statistical analysis, providing a measure of total error/adaptation during the task.

Statistical analyses were conducted in MATLAB, R (R Core Team, version 3.6.3) and SPSS (IBM, version 26). All ANOVAs were run in general linear model format and followed up with Bonferroni-corrected multiple comparisons, when a significant main effect or interaction was found. The threshold for statistical significance was set at p < 0.05 and we report partial eta squared (ηp^2^) effect sizes for ANOVAs. Estimation statistics were conducted as per [27], reporting paired Cohen’s d effect sizes and bias-corrected and accelerated confidence intervals (following 5000 bootstrap samples).

Data from the secondary experimental group were processed and analysed separately as they were collected after the main experiment and this group was not included in the original study design.

## Results and Discussion

We first sought to determine any differences in adaptation between trial contexts for each stimulation group during the task. We measured the area under the learning curve (calculated from lateral deviation of movements from the target midline; see methods for further details) in order to compare adaptation performance on trials performed with simultaneous stimulation and those without, using a 3×3×2 mixed-design ANOVA. The ANOVA contained within-subject factors of adaptation phase (baseline, adaptation, washout) and trial context (CW (left-shift), CCW (right-shift)) and the between-subject factor of stimulation group (M1 er-TDCS, cerebellar er-TDCS, sham). The ANOVA revealed a significant main effect of adaptation phase (F (2, 342) = 1312.45, p < 0.001, η^2^p = 0.89), stimulation group (F (2, 342) = 18.93, p < 0.001, η^2^p = 0.1), and a significant three-way interaction between group, phase and trial context (F (4,342) = 3.44, p = 0.009, η^2^p = 0.04). Bonferroni corrected multiple comparisons revealed that, during adaptation, participants who received cerebellar er-TDCS made significantly less error, and thus adapted better, on stimulated CCW trials compared to unstimulated CW trials, p < 0.001 (Figure 2). In contrast, we found no evidence of enhanced adaptation following M1 er-TDCS, as participants made similar levels of error on both stimulated and unstimulated trials (p = 0.97). Similarly, participants in the sham group adapted comparably during left and right-shift trials (p = 0.72), suggesting there was no in-built task bias (Figure 2).

**Figure 2:**
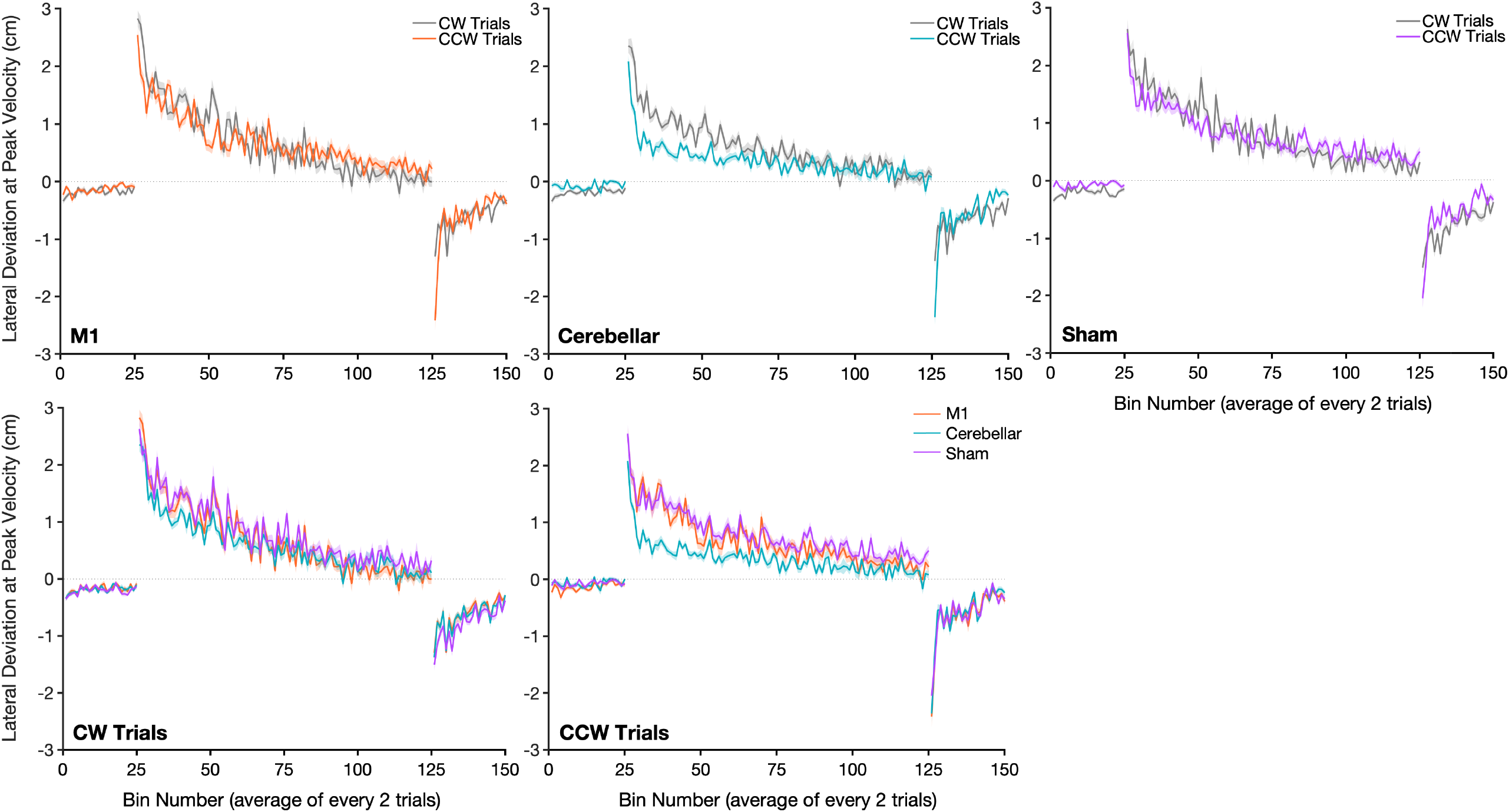
Cerebellar er-TDCS selectively improves context-dependent force-field adaptation. **Top panel:** Mean lateral deviation for CW (left-shift) and CCW (right-shift) trials (± standard error, shaded regions), averaged into bins of two trials for the M1 er-TDCS, cerebellar er-TDCS and sham stimulation groups. The M1 and cerebellar groups received er-TDCS on CCW trials during the adaptation phase. **Lower panel:** mean LD (bins of two trials) for all three stimulation groups during either CW or CCW contextual trials.

Testing the robustness of these effects using estimation statistics [27], confirmed that er-TDCS applied to the cerebellum selectively improved the adaptation on CCW trials, while CW trials were unaffected. The effect size for this comparison was substantial (paired Cohen’s d = -1.3) with 95.0% confidence intervals (CI) that did not overlap zero ([-2.05, -0.53]; Figure 3). Conversely, the paired Cohen’s d between CW and CCW trials for the M1 er-TDCS group was close to zero, which fell well within the CI (d = -0.0059, 95.0% CI = [-0.66, 0.64]); for the sham group d = -0.061 and 95.0% CI = [-0.69, 0.57].

**Figure 3:**
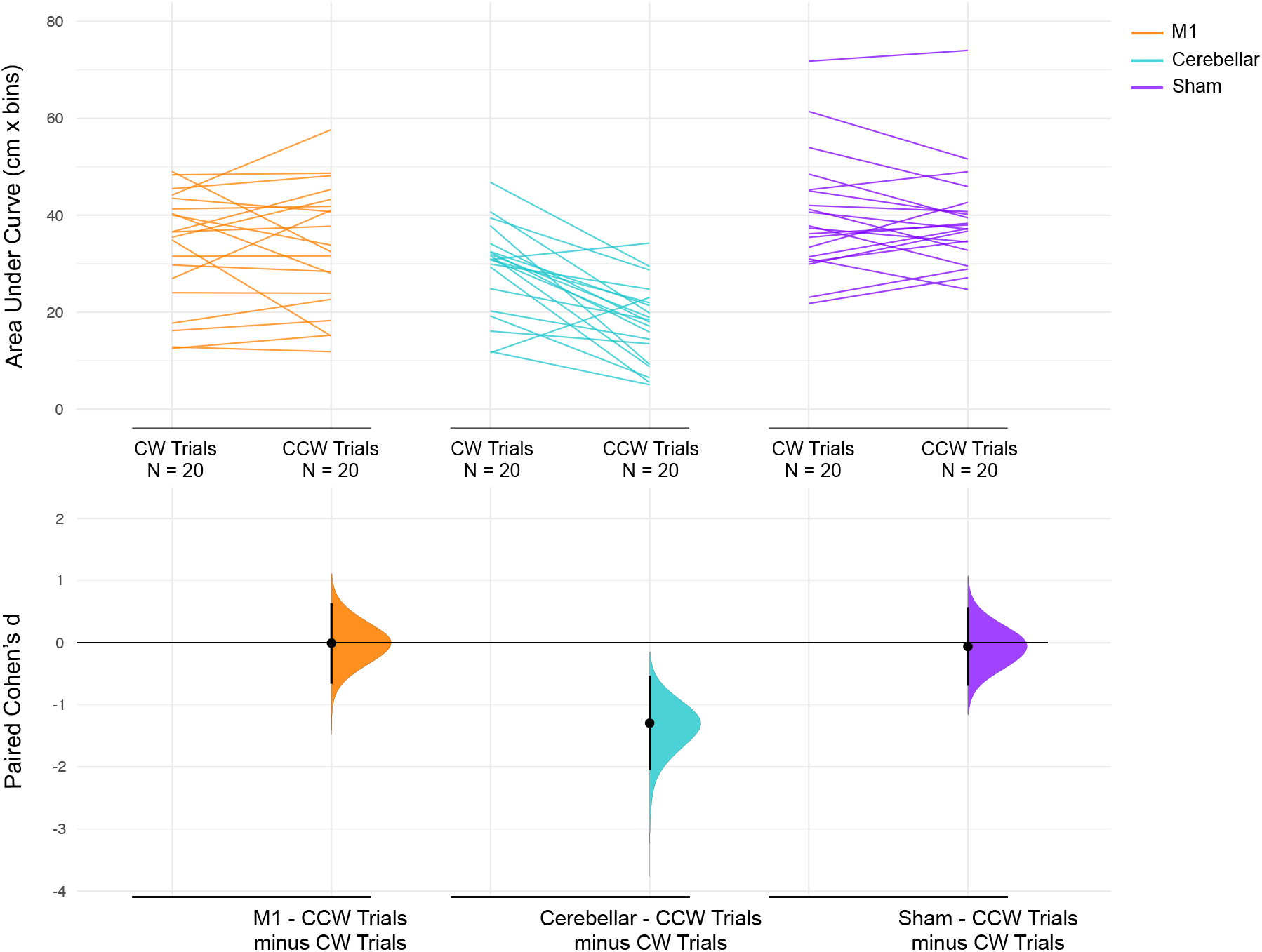
Direct comparisons of adaptation during CW and CCW force-fields. **Top panel:** area under the curve (cm x bins) for CW vs CCW trials for each participant during the adaptation phase, with each participant’s data connected by a line. **Lower panel:** paired Cohen’s d plotted as a bootstrap sampling distribution. Mean differences are depicted as dots, with 95.0% CIs indicated by black vertical bars.

Further multiple comparisons from the three-way interaction suggest that there no differences in adaptive performance between trial contexts during baseline for any of the stimulation groups (all p > 0.504), indicating that any differences during adaptation were not a result of uneven performance during baseline (Figure 2).

There were also no differences in de-adaptation between trial contexts during washout trials (all p > 0.23), which is likely a result of similar levels of learning reached towards the end of the adaptation phase. Additionally, when considering overall adaptation (adaptation to both CW and CCW trials, sign transformed and combined), results from the ANOVA revealed that participants in the cerebellar er-TDCS group made significantly less error compared to the M1 er-TDCS and sham group (both p < 0.001), with no significant differences between the latter two groups, p = 0.078.

These results show that event-related stimulation of the cerebellum selectively enhanced adaptation of stimulated CCW trials compared to CW trials, with no effect of M1 er-TDCS on context-dependent adaptation or task bias (indicated by sham data). The reduction of error on CCW (right-shift) trials following cerebellar er-TDCS also contributed to (at least in part) an overall improvement in adaptation during the task (Figure 2 & Supplementary Figure 1).

Results from our additional experimental group helped to confirm the findings from our main experiment. We found that er-TDCS over the cerebellum enhanced adaptation of movements through the stimulated context compared to the unstimulated control – a replication of the effect initially found (see Figure 4a). Area under the learning curve was compared in a 3×2 way ANOVA (Task Phase: baseline, adaptation, washout; Trial Context: CW (left-shift), CCW (right-shift)), and revealed significant main effects of task phase (F (2,102) = 248.23, p < 0.001, η^2^p = 0.83), trial context (F (1,102) = 12.49, p < 0.001, η^2^p = 0.11) and a significant interaction (F (2,102) = 6.05, p = 0.003, η^2^p = 0.11). Bonferroni corrected multiple comparisons showed that during the adaptation phase participants made significantly less error on stimulated CW trials, compared to CCW trials (p < 0.001). We found no evidence to suggest performance during the two trial contexts were different during baseline or washout phases (both p > 0.53). Estimation statistics affirmed this result and revealed a large effect size (paired cohen’s d = 0.82, 95% CI = [0.095, 1.43]), with zero falling outside the 95% CI range (see Figure 4b).

**Figure 4:**
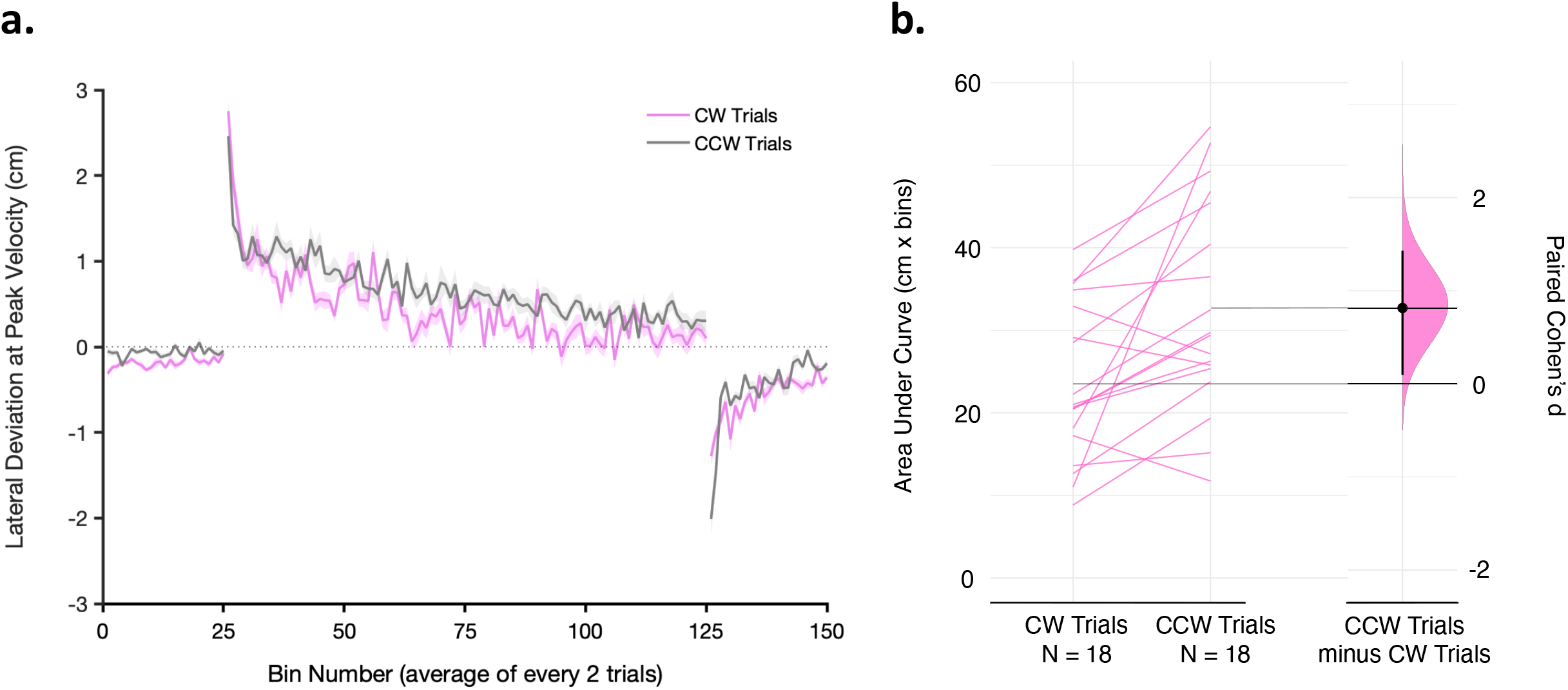
Adaptation performance and context selectivity for the secondary experimental group receiving er-TDCS during CW trials. **a:** Mean lateral deviation for CW (left-shift) and CCW (right-shift) trials (± standard error, shaded regions), averaged into bins of two trials for the secondary experimental group. Participants in this group received er-TDCS over the cerebellum during CW adaptation trials. Panel **(b.)** depicts the area under the curve (cm x bins) for CW vs CCW trials for each participant during the adaptation phase, with each participant’s data connected by a line in a Paired Gardner-Altman plot, with the paired Cohen’s d plotted as a bootstrap sampling distribution alongside. The mean difference is shown as a dot, with 95.0% CIs indicated by black vertical bars.

One concern was that short epochs of stimulation, with rapid onset/offset, might be perceived by the participant and act as a form of attentional cue. However, we found no significant differences between their self-reported Confidence in Stimulation (F (3,74) = 0.078, p = 0.97, η^2^p = 0.003), suggesting equivalent blinding to the stimulation condition was achieved. Given that this er-TDCS protocol is relatively novel, it is also pleasing that the short bouts of TDCS did not cause excessive discomfort (Perceived Comfort: F (3,74) = 2.31, p = 0.084, η^2^p = 0.085), with all groups reporting low levels of discomfort. Additionally, comfort levels were similar to those reported after 17 minutes of continuous TDCS [21]). Crucially, only one participant (in the M1 er-TDCS group) noticed that the stimulation only occurred on CCW force-field trials with a corresponding right-shift in task display. The lack of awareness of stimulation timing in the cerebellar group suggests enhanced adaptation cannot be due to explicit cueing or other explicit mechanism such as increased attention towards CCW (right-shift) trials.

These initial results suggest that brief periods of TDCS applied over the cerebellum and in synchrony with movement can selectively and specifically improve motor adaptation of that movement, while leaving adaptation of interleaved movements unaffected. Although many studies have reported positive effects of TDCS on motor learning and rehabilitation, when applied continuously for 10-20 minutes [21, 28–34], there are a growing number of studies reporting null or mixed effects [35–38], leading to uncertainty around the effectiveness of TDCS [39]. During continuous stimulation, for 10-20 minutes, any number of different behaviours may be performed alongside the specific task that is the ‘target’ of stimulation. The concatenation of all these behaviours under the same stimulation conditions may lead to changes in excitation levels in multiple cortical circuits that confound the results, contributing to some of the conflicting findings [40]. In contrast, event-related TDCS may selectively modulate only those circuits and task-related synapses that are contemporaneously active and undergoing concurrent plasticity [16, 41]. As such, er-TDCS could prove beneficial when long experimental or rehabilitative protocols are required, as recent research suggests that long continuous bouts of TDCS may cause the resultant modulatory effects to diminish and even reverse over time, i.e., from excitation to inhibition [42]. Neural recording studies have also found that the effects of direct current stimulation may attenuate during long stimulation blocks. These changes have been seen in spontaneous neural activity and evoked potentials, due to short-term habituation-like adaptation processes [43–45].

The plastic mechanisms responsible for the improvement in adaptation of specific movements associated with er-TDCS in this study are still unclear. We propose that (1) applying anodal TDCS concurrently during movement enhances activity within the neural circuit associated with and activated by this specific behaviour [16]. This increased activity in turn could potentiate the synapses within this circuit that are ‘eligible’ for Hebbian change during the stimulated behaviour. (2) There are homeostatic mechanisms that act to down-regulate the increase in activity [43-45], that might be of slow onset, most prominent during prolonged TDCS. (3) By providing brief er-TDCS, it is more likely that only those circuits involved in the particular (concurrent) behavioural context are the ones that are potentiated, without homeostatic reduction. In essence, er-TDCS provides transient heightened potentiation that ‘focuses’ Hebbian learning onto specific neurobehavioural circuits [41, 46].

However, there are other plausible mechanisms that are less reliant on synaptic plasticity mechanisms. For instance, by boosting the excitability of circuits of cerebellar neurons during movement, er-TDCS may act to increase the number of active neurons within the circuit associated with the specific behaviour. This would result in a greater network of cerebellar neurons selectively activated during adaptive movements, which in turn, could plausibly drive better, more accurate movements and result in improved adaptation trial-by-trial.

It is also important to consider which learning processes which may be responsible for this enhanced adaptation. Given that stimulation was ramped up for 1 second prior to movement initiation, and outlasted the movement, there is scope to suggest that activity related to motor planning and preparation may have been facilitated before each stimulated trial, or that error processing after each trial was facilitated. Additionally, as sensorimotor adaptation is highly dependent on the cerebellum [47– 49], it is possible that er-TDCS modulates forward model processing, leading to enhanced prediction error and thus more rapid learning during the stimulated context. Experiments testing some of these suggestions will be reported in due course.

We should also note that there is some non-uniformity in the apparent mass of the vBOT when moving in different directions due to the direction of the ‘elbow’ joint (Figure 1a). This may have resulted in a small disparity in the errors between the two movement conditions when the CW and CCW forces were first introduced, as seen for example in the sham group data (Figure 2). However, this difference was quickly overcome in the sham and M1 group, and we found significantly improved adaptation in both CW and CCW cerebellar conditions, so it is unlikely to have had major impact on the overall results.

Hebbian-like plasticity can also be induced in the motor cortex, for example using paired associative stimulation (PAS) and use-dependent plasticity (UDP) protocols [50–53]. However, we found no specific effect of M1 er-TDCS. Adaptation of CW and CCW trials were not significantly different, and participants performed similarly to the sham stimulation group. The null effect of M1 er-TDCS may be explained by the nature of the task. Reach adaptation to force-fields is believed to be predominantly cerebellar-dependant [48, 53–56], and is dominated by the proximal muscles of the arm [21, 26]. Thus, targeting M1 may have had relatively little effect on the processes governing motor adaptation/performance during the task.

The results presented here suggest ways in which TDCS could be utilised in research and rehabilitation, with a focus on increased temporal resolution. For example, er-TDCS could be used in conjunction with therapy protocols comprising of precise temporal epochs, with further scope beyond the motor modality e.g., targeting memory encoding or recall [57, 58]. While TMS remains a more temporally precise non-invasive brain stimulation intervention, er-TDCS may now be considered, especially given that direct current stimulation has the potential to modulate neural excitability bi-directionally. It is yet to be determined how this novel stimulation protocol will transfer to other settings, and future research in diverse learning contexts, targeting different brain regions, will be required to determine its robustness.

In conclusion, we have shown that brief epochs of TDCS over the cerebellum can selectively modulate motor adaptation when delivered coincidentally with reaching movements. We suggest that the coupling of stimulation and movement influences mechanisms of Hebbian-like plasticity and thus facilitates learning in the stimulated context. These initial results potentially open up new possibilities for the use of TDCS in both research and clinical settings, to improve its effectiveness and specificity. They also demonstrate that stimulation can be applied in very short bouts without inducing severe discomfort or side-effects, and without the participants’ explicit knowledge of the specific stimulation protocol.

## Supporting information

Supplementary Information & Analysis

## Acknowledgements

This work was supported by the MRC-Versus Arthritis Centre for Musculoskeletal Ageing Research (CMAR). We thank an anonymous reviewer for their helpful suggestions for the discussion of our results.

## Author Contributions

**Matthew Weightman:** Conceptualization, Methodology, Investigation, Formal analysis, Visualization, Writing - original draft. **John-Stuart Brittain:** Conceptualization, Methodology, Formal analysis, Writing - review & editing. **Alison Hall:** Methodology, Writing - review & editing. **R. Chris Miall:** Conceptualization, Formal analysis, Writing - review & editing.

**Ned Jenkinson:** Conceptualization, Methodology, Formal analysis, Writing - review & editing.

## Competing Interests

The authors declare no competing interests exist

## Data and Code Availability

Data used in this manuscript to create figures have been deposited to Mendeley Data and are available at:

DOI: 10.17632/2k33svf244.2

For further information and requests for resources please communicate with corresponding author, Matthew Weightman.

## Notes

https://doi.org/10.17632/2k33svf244.2

### Competing Interest Statement

The authors have declared no competing interest.

