## Supplementary Information & Analysis for "Timing is everything: event-related transcranial direct current stimulation improves motor adaptation"

**Affiliations:**

The University of Birmingham

Edgbaston

Birmingham

B15 2TT

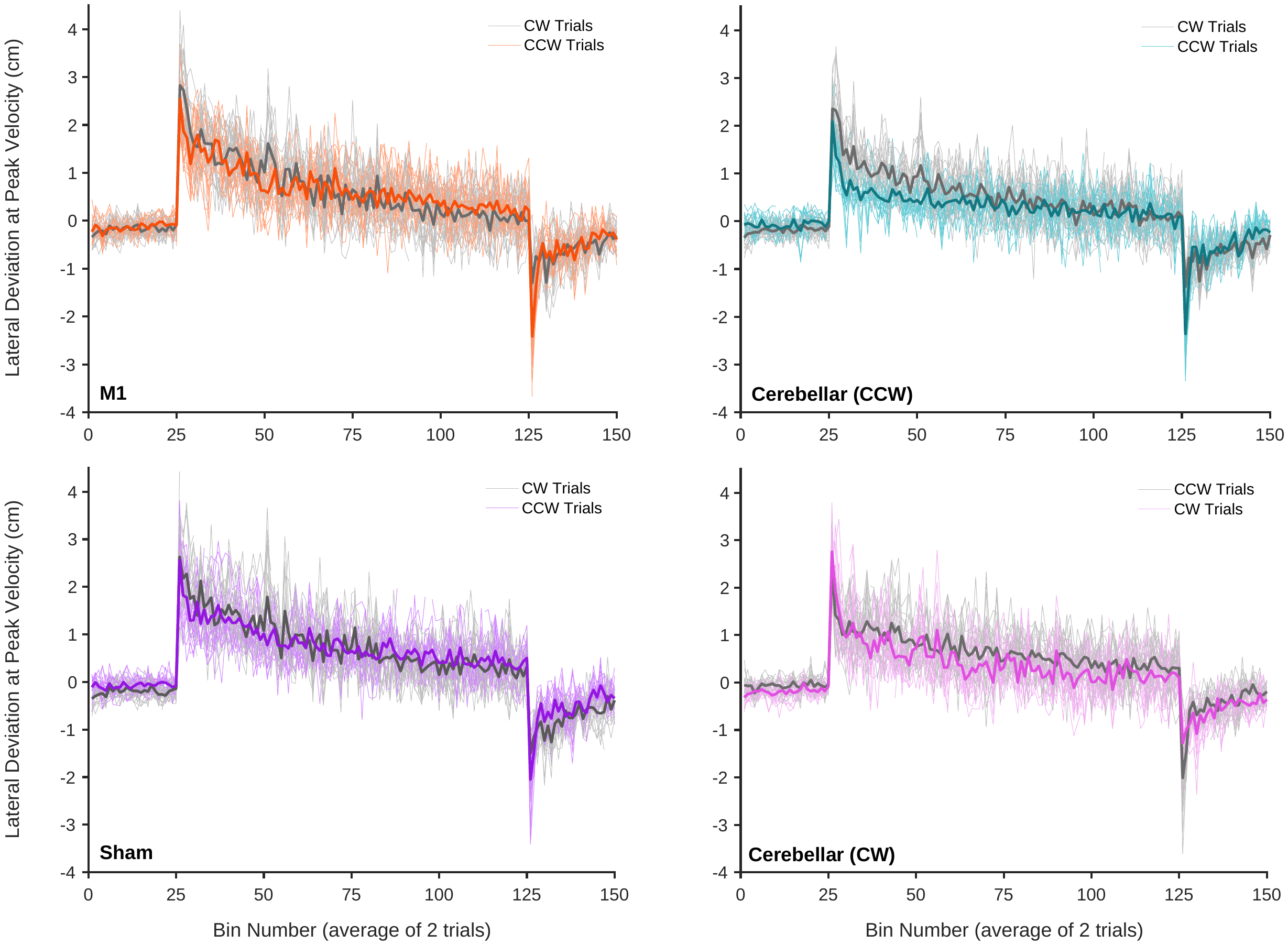
**Supplementary Figures**

**Supplementary Figure 1:** Individual data traces for lateral deviation.

*Lateral deviation for CW (left-shift) and CCW (right-shift) trials averaged into bins of two trials for the M1 er-TDCS, cerebellar er-TDCS, sham stimulation and the secondary cerebellar er-TDCS groups. The M1 and cerebellar groups received er-TDCS on CCW trials during the adaptation phase and the secondary experimental group (cerebellar (CW)) received er-TDCS during the adaptation phase on CW trials. Individual traces are plotted for each participant for each trial context, with the group mean superimposed in bold on top.*

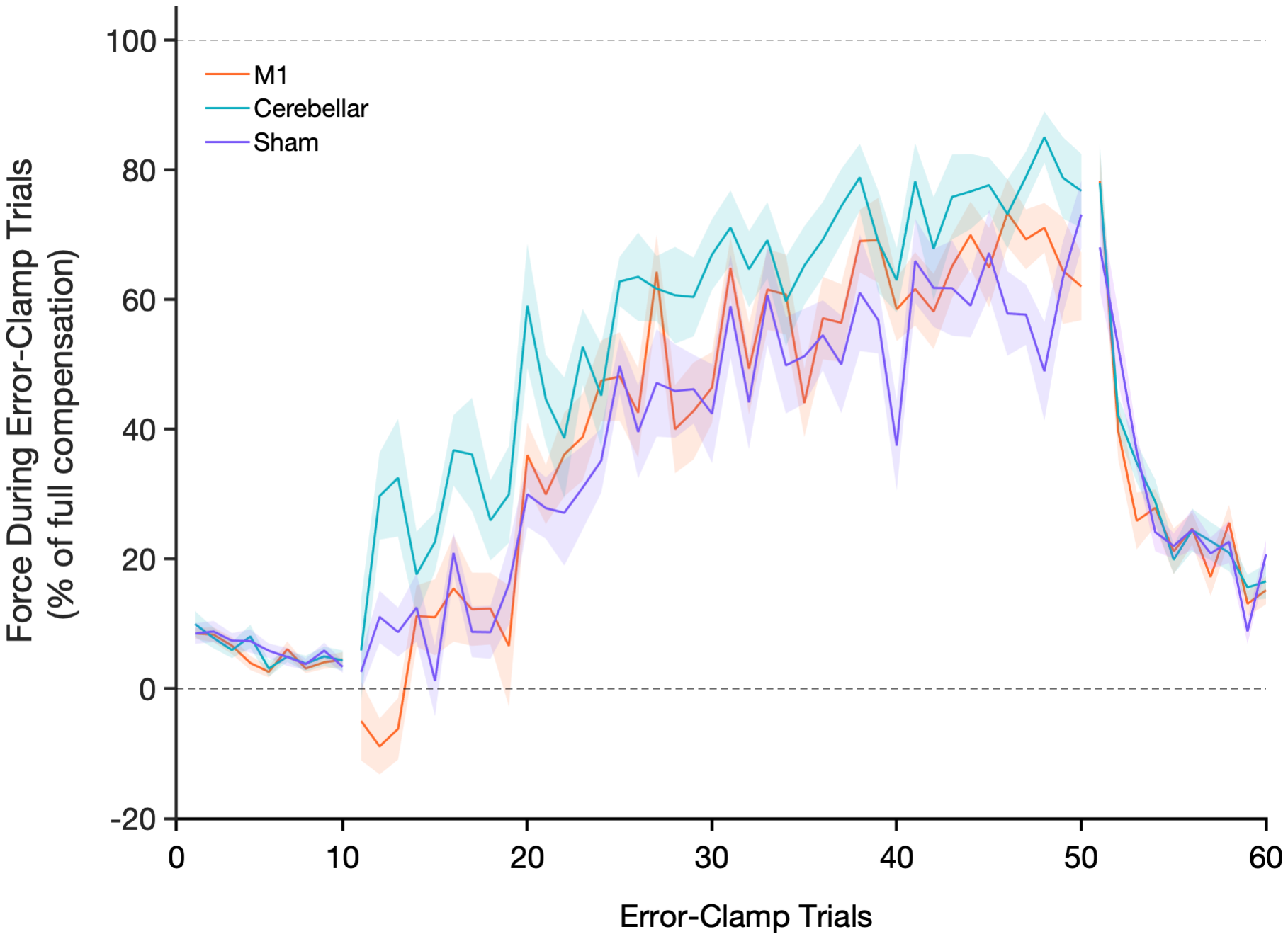
**Supplementary Figure 2:** Force compensation during error-clamp trials, main experiment.
*Average force compensation during error-clamp trials (±standard error, shaded region) in each task phase for all three stimulation groups. Data is sign adjusted to show the percentage of full compensation for both left and right-shift error-clamp trials. Area under the curve was calculated (AUC-compensation) for each participant in each task phase and trial context, in order to compare performance. A 3x3x2 (task phase x stimulation group x contextual shift) mixed-design ANOVA revealed significant main effects of task phase (F (2, 342) = 873.68, p < 0.001, ηp^2^ = 0.84) and stimulation group (F (2, 343) = 10.78, p < 0.001, ηp^2^ = 0.06), but no significant three-way complex interaction (F (4, 342) = 0.66, p = 0.99, ηp2 = 0.001). There were no differences between force compensation on left vs right-shift trials for any of the stimulation groups, all p > 0.072, suggesting er-TDCS had no specific effect on performance during error-clamp trials. The ANOVA, however, did reveal a significant interaction between phase and stimulation group (F (4, 342) = 9.92, p < 0.001, ηp^2^ = 0.1). Multiple comparisons showed that the three stimulation groups performed similarly on error-clamp trials during baseline and washout phases (all p > 0.99), yet during the adaptation phase, participants in the cerebellar er-TDCS produced greater levels of force compensation compared to the M1 er-TDCS and sham group, both p < 0.001, with no difference between the latter two group (p = 0.39). These results may suggest that er-TDCS over the cerebellum had a global effect on force compensation during error-clamp trials,* *rather than the specific timing-dependent effect observed for lateral deviation error-reduction.*

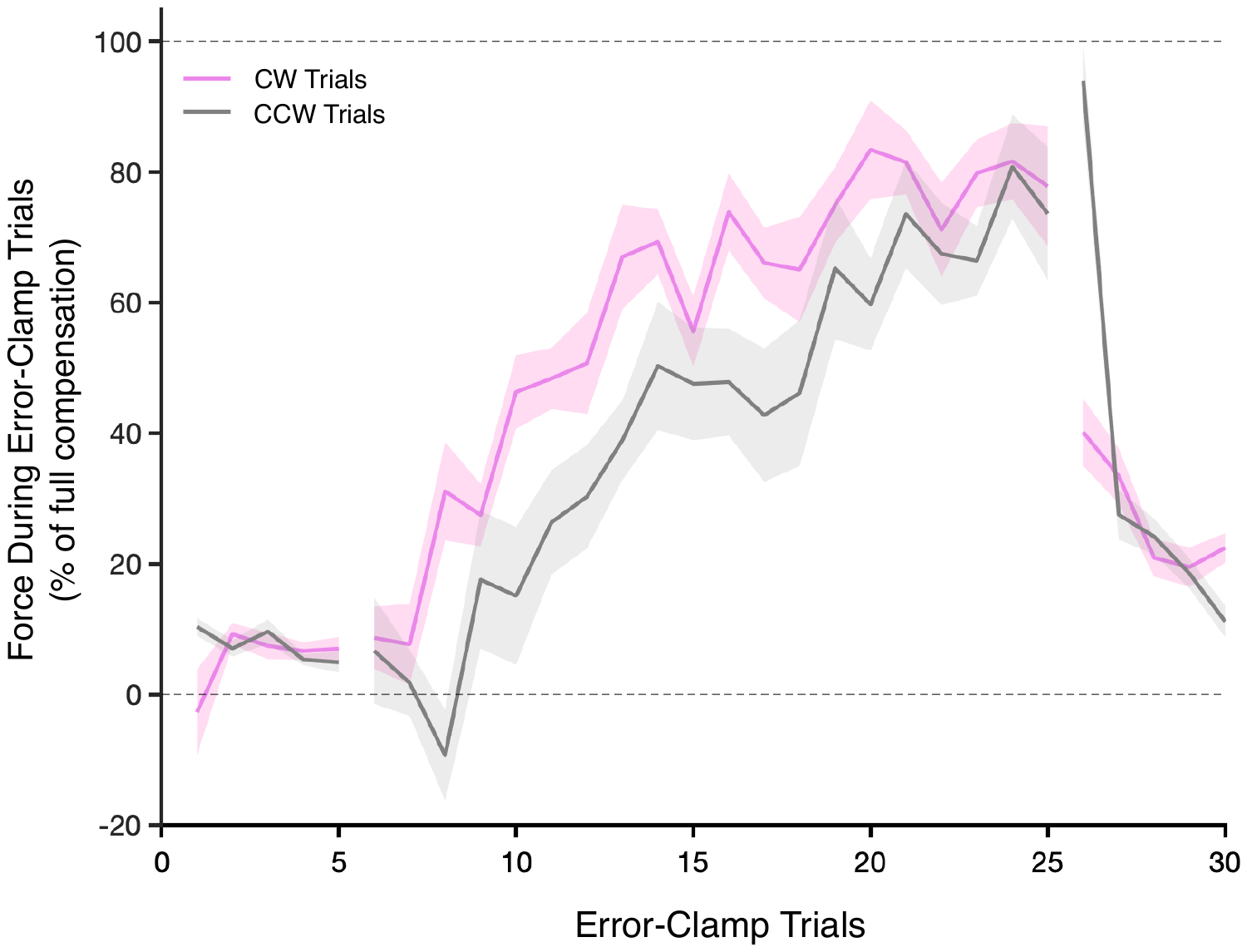
**Supplementary Figure 3:** Force compensation during error-clamp trials, secondary experimental group.

*Average force compensation during CW and CCW error-clamp trials (± standard error, shaded region) for data from the secondary experimental group. For this group, cerebellar er-TDCS was applied on CW (right- shift) error-clamp trials (trials 6-25 on figure). Data is sign adjusted to show the percentage of full compensation for both left and right-shift error-clamp trials.
Specific enhancement in force compensation was found during error-clamp trials in the stimulated context (CW). Thus, participants applied more force against the channel wall during expected CW adaptation trials than CCW trials. Area under the curve during error-clamp trials were compared in a 3x2 way ANOVA (Task Phase x Trial Context) and revealed significant main effects for Phase (F(2, 102) = 192.26, p < 0.001, ηp^2^ = 0.79), Context (F(1, 102) = 4.24, p = 0.042, ηp^2^ = 0.04) and a significant interaction between trial phase and context (F(2, 102) = 5.7, p = 0.005, ηp^2^ = 0.1). Comparisons from this interaction showed greater compensation on stimulated CW compared to CCW trials (p < 0.001) during the adaptation phase, with no differences for baseline and washout trials (all p > 0.83).*
